# Mitochondrial Dysfunction in Endothelial Cells Drives Greater Vascular Impairment in Females with Diabetes-Associated Peripheral Artery Disease

**DOI:** 10.1101/2025.09.22.677648

**Authors:** Siân P Cartland, Elaina Kelland, Natalie Le, Jaideep Singh, Madeleine Murphy, Rachael Menezes, Jacqueline Ku, Lorna Beattie, Lisa Turner, Jacky Loa, Lauren Boccanfuso, Lauren Sandeman, Millie Jiang, Polina E Nedoboy, Malathi I Dona, Sergey Tumanov, Joseph E Powell, Alexander R Pinto, Shane R Thomas, Christina Bursill, David Celermajer, Christopher P Stanley, Sarah J Aitken, David A Robinson, Mary M Kavurma

## Abstract

**Background:** Women with peripheral artery disease (PAD) experience poorer clinical outcomes than men, particularly in the setting of diabetes. However, the mechanistic basis for these sex-specific disparities remains unclear.

**Methods:** Here, we investigated endothelial cell (EC) function(s) in diabetes-associated PAD, with a focus on sex differences. Limb tissues from patients with diabetes and chronic limb-threatening ischemia (CLTI) undergoing amputation, and a diabetes mouse model of hindlimb ischemia (HLI), were assessed for vasodilatory capacity, angiogenesis, oxidative stress and changes to expression of mitochondrial complex genes. ECs exposed to a hyperglycemic environment *in vitro* were assessed for mitochondrial function. The therapeutic potential of the mitochondrial-targeted antioxidant MitoQ was investigated.

**Results:** ECs from females with diabetes-associated PAD have altered responses compared to males. Specifically, limb vessels and skeletal muscle from females exhibit reduced arterial relaxation, angiogenesis and increased oxidative stress in response to HLI in mice, and in tissues from patients. Single-cell RNA sequencing of murine limbs revealed marked suppression of EC mitochondrial complex genes in females with diabetes. Female human ECs exposed to high glucose had reduced respiration, reduced expression of mitochondrial genes and increased oxidative stress. Remarkably, MitoQ restored arterial relaxation and the angiogenic response in female diabetes-associated PAD.

**Conclusion:** Our findings uncover a striking sex-specific vulnerability involving oxidative stress and mitochondrial dysfunction in EC health in diabetes-associated PAD. These results highlight the need for sex-specific therapeutic strategies in diabetic PAD, which might include mitochondrial targeted antioxidant strategies.

## INTRODUCTION

Peripheral artery disease (PAD) in the context of diabetes is an increasingly urgent problem with distinct clinical challenges^1^. Approximately 20% of individuals with diabetes have PAD^2^, a disease burden that is expected to grow substantially as global diabetes prevalence is projected to rise by over 50% by 2049^3^. Diabetes not only accelerates the onset and severity of PAD, but also modifies its underlying vascular biology, promoting endothelial dysfunction, inflammation, and impaired angiogenesis. Importantly, while women and men have a similar overall prevalence of PAD, women with diabetes appear more susceptible to severe PAD, often presenting with more advanced ischemic manifestations^4^. This contributes to an increased rate of adverse treatment outcomes observed in women compared to men, including more complications during surgical or endovascular revascularisation^5,6^, early graft thrombosis, amputation and mortality^1,7^. These differences underscore a critical need for a greater understanding of the mechanistic pathways by which diabetes contributes to atherosclerosis and how sex-specific factors shape disease development.

Endothelial dysfunction is a hallmark of PAD^8^, a condition in which the endothelium loses its ability to promote vasodilation, stimulate new blood vessels and suppress oxidative stress and inflammation. While these features are well recognized, much less is known about how diabetes might modify endothelial cell (EC) behavior in a sex-specific manner. A recent study investigating sex differences in atherosclerosis identified gene clusters that were predominantly expressed by the female endothelium^9^, highlighting the endothelium as a key target for understanding biological sex differences. Extensive clinical and experimental evidence indicates that diabetes is a major driver of EC dysfunction, impairing nitric oxide bioavailability, disrupting redox homeostasis, and weakening the protective barrier function of endothelial cells (reviewed in^10–13^). Sex-based mitochondrial differences have been identified^14^, with female ECs showing reduced respiratory activity, impaired angiogenic potential, and elevated reactive oxygen species (ROS) compared to male ECs^15^. However, the extent to which these mitochondrial and endothelial disparities contribute to diabetes-associated PAD remains poorly defined.

To address this gap, we examined how diabetes influences endothelial function in a sex-specific manner. Our findings demonstrate that female PAD patients, diabetic mice, and cultured human ECs exhibit more pronounced impairments in vasodilation, angiogenesis and mitochondrial activity compared to their male counterparts. These observations reinforce the need for therapeutic strategies that target mitochondrial dysfunction as a central modulator of endothelial bioenergetics, redox balance, and vascular responsiveness. In this context, we tested MitoQ, a mitochondria-targeted antioxidant, and found that it restored vasodilatory and angiogenic responses in female diabetic mice and reduced oxidative stress in ECs. These results suggest that improving mitochondrial function may offer a promising avenue to enhance endothelial health and microvascular recovery, particularly in women with diabetes-associated PAD, and may support better outcomes following revascularization or surgical interventions.

## CLINICAL PERSPECTIVE

### What is new?

- Female patients with diabetes-associated PAD patients exhibit reduced vasodilation and lower microvascular density in skeletal limb muscles when compared to male patients, with more localized oxidative stress.
- In both *in vitro* and *in vivo* models, diabetes impairs endothelial mitochondrial function more in females than males, resulting in alterations to cell function that compromise the endothelium’s homeostatic responses e.g., female mice had reduced vasodilation, angiogenesis and wound healing but elevated oxidative stress.
- Treatment with the mitochondrial-targeted antioxidant MitoQ, improved endothelial function in female diabetic mice, restoring angiogenic and vasodilatory responses, in part by reducing mitochondrial oxidative stress.

### What are the clinical implications?

Women with diabetes and PAD experience more severe clinical ischemic complications than men, which appear to be partly driven by sex-specific endothelial and mitochondrial dysfunction. Targeting endothelial mitochondrial impairment with MitoQ—a clinically available mitochondrial antioxidant—may offer a therapeutic strategy to reduce ischaemic damage from PAD in female patients with diabetes.

## METHODS

### Human studies

Male and female diabetic patients aged 18 years and older presenting with chronic limb-threatening ischemia (CLTI) and requiring above- or below-knee amputation at Royal Prince Alfred Hospital in Sydney, Australia, were enrolled in the study. Blood plasma was collected, snap-frozen, and stored at -80°C. Skeletal muscle samples were obtained from the tibialis anterior, near the amputation site. The tissue was split, fixed in formalin for histology or flash frozen for gene expression studies. For myography, arteries were harvested from the subcutaneous fat surrounding the tibialis anterior, near the cut site as previously described by us^16^. The study was approved by the Sydney Local Health District (SLHD) Human Ethics Committee (X20-0183; Sydney, Australia) and all patients provided written informed consent.

### Mice

*Apoe^-/-^* mice on a C57Bl6 background were bred at Australian BioResources (ABR; Mossvale, NSW, Australia) or purchased from Ozgene Australian Resource Centre. C57Bl6 mice were purchased from ABR. All mice were housed with 12:12 h light-dark cycles with free access to food and water. Mice were anaesthetized by inhalation of 2-3% isoflurane prior to euthanasia via cardiac exsanguination. A mix of male and female mice were used in all experiments. These studies were approved by the SLHD Animal Welfare Committee (Sydney, Australia, 2020-008 and 2024-007) and Institutional Biosafety Committee (Sydney Australia, 18-021 and 23-027) in accordance with the Australian code for the care and use of animals for scientific purposes (National Health and Medical Research Council, NHMRC).

### Diabetes-associated PAD model

A schematic of the model is described in Supplementary Figure 1. Male and female *Apoe^-/-^* mice, ∼8-10 weeks of age, were acclimatised for 1 w followed by the induction of diabetes using streptozotocin (STZ; Sigma-Aldrich, Sydney, NSW, Australia) at 75 mg/kg i.p., for 5 consecutive days. Vehicle, citrate buffer served as the non-diabetic control. Females were given a second boost of STZ at 75 mg/kg for 5 consecutive days at week 11 after the initial STZ dose. Animals remained on standard chow. Weight and unfasted blood glucose were measured weekly for 20 weeks. Animals with non-fasted blood glucose <15mmol/L were not considered diabetic and removed from the study. Fifteen weeks after the initial diabetes induction, a glucose tolerance test was performed; 1 g/kg body weight D-glucose (Sigma-Aldrich, Sydney, NSW, Australia) by i.p., following overnight fasting. Plasma glucose levels were measured using a glucometer (LifeSmart, Genesis Biotech Pty Ltd; Brisbane, Qld, Australia) with blood collected by tail vein nick. Eighteen weeks after the initial induction of diabetes, hindlimb ischemia (HLI) surgery was performed^16–19^. Mice were anesthetized and fur removed from the surgical site, which was then cleaned with ethanol. The femoral artery and vein in the left hindlimb were exposed, separated from the nerve, ligated, and excised. The right hindlimb was used as the control, non-ischemic limb. Limb blood perfusion was evaluated by Laser Doppler Imaging (moorLDI2-IR; Moor Instruments, Axminster, United Kingdom) prior to surgery and at day 1, 3, 7 and 14 post-surgeries. The laser Doppler perfusion index (LDPI) was calculated as the ratio of the ischemic:non-ischemic limb for each mouse. Fourteen days later mice were euthanized. Blood was collected for plasma, which was snap-frozen and stored at -80°C. The gastrocnemius muscle was isolated and cut in half lengthways, one half formalin-fixed for immunohistochemistry (IHC), and the other half digested for single cell RNA (scRNA-seq) sequencing, or flash frozen. The aortae and soleus muscle were also isolated, and flash frozen for gene expression studies. Brachiocephalic arteries were isolated and fixed in formalin to assess extent of atherosclerosis^20–22^.

### MitoQ treatment

Adult male and female *Apoe^-/-^* mice were rendered diabetic using STZ at 75 mg/kg i.p., for 5 consecutive days. Ten to fourteen days later, mice underwent HLI. During surgery, Alzet® Osmotic Pumps were inserted subcutaneously in the backs of mice, delivering 10 µg of MitoQ (Selleckchem; Houston, TX, USA) per day for 14 days. Blood perfusion was measured using Laser Doppler Imaging and mice were euthanized 14-days-post HLI. Indicated tissues were collected.

### Wound healing

*Apoe^-/-^* male and female mice aged 8-11 weeks were rendered diabetic with STZ by i.p. (165 µg/g, males or 180 µg/g for females). Blood glucose was measured weekly. Blood glucose <15mmol/L were not considered diabetic and removed from the study. The wound healing surgery was performed as previously described^23^. Briefly, under isoflurane anaesthesia two 5 mm diameter full thickness wounds were created on the back flanks and secured with silicon splints, attached with superglue and sutures. Each wound received 10 µl PBS topically and was then covered with a dressing (Opsite^TM^) daily. Wound areas were measured daily and wound blood perfusion determined by Laser Doppler Imaging. Ten days post-surgery, mice were euthanized. Each wound was excised: one-half flash frozen for gene expression, the other half fixed in 20% sucrose/PBS for 24 h, 4% paraformaldehyde/PBS for 24 h, then 70% ethanol at 4°C and processed for histology. This procedure was conducted with ethical approval from the South Australian Health and Medical Research Institute Animal Ethics Committee (SAM20.005).

### Myography

Myography was performed on isolated mouse and human arteries^16^. Briefly, dissected arteries were cleaned, sectioned into 2-mm segments and mounted on tungsten wires (40 µm diameter) on a myograph system (Danish Myo Technology) at 37°C. Arteries were maintained in Krebs-Henseleit solution and gassed with 5% CO[ in O_2_. Isometric tension was recorded using force displacement transducers and analyzed with LabChart V8Pro (ADInstruments). Vessels were normalized to an internal diameter corresponding to 90% of 13.3 kPa. Following normalization, arteries were contracted using 120 mM KCl until a stable contraction was achieved, subsequently washed, rested for 30 min, and assessed for endothelium-dependent or independent relaxation using increasing doses of acetylcholine or sodium nitroprusside.

### Plasma cholesterol

Total plasma cholesterol was measured using the Cholesterol E commercial kit (Wako Chemical, Osaka, Japan, 439-17501), according to manufacturer instructions.

### Immunohistochemistry

Histology was performed on 3 µm (brachiocephalic artery) or 5 µm (all other tissue) sections from indicated mouse and human tissues. Hematoxylin and eosin (H+E) were used to assess tissue architecture. Myocyte area was assessed from H+E sections using ImageJ. Five regions of interest were randomly selected per sample and area of 50 myocytes measured per region of interest. Immunofluorescence was performed to quantify the number of microvessels or CD31 density using anti-CD31 (1:100; AF3628, In Vitro Technologies), anti-SMA (1:500; F3777, Sigma Aldrich) and/or anti-laminin (1:250; NB300-144, In Vitro Technologies) antibodies in mice (gastrocnemius, wounds) or human tissues (tibialis anterior). Non-conjugated primary antibodies were detected using anti-IgG Cy5 (1:500, Jackson ImmunoResearch, 711-175-152) and Cy3 (1:500, Jackson ImmunoResearch, 705-166-147) fluorophores. Slides were mounted with DAPI mounting media (SouthernBiotech, 0100-20). Control sections excluding primary or secondary antibodies were negative. Images were captured using a Zeiss Axio Imager Z2 or Zeiss Axio Scan.Z1 microscope. CD31+SMA+ microvessels <50 µm were counted and normalized to myocyte number within regions of interest using Zen 3.6 software (Zeiss). For mouse studies, numbers of CD31^+^SMA^+^ microvessels per myocyte number in the ischemic limb were normalized to numbers in the control limb. For wounds, CD31 density was assessed^24^. Myocyte area from human muscles was measured in the tibialis anterior using image J^21^, assessing the area of 50 myocytes in 3-5 regions of interest per patient.

## 8-Isoprostane ELISA

The 8-isoprostane ELISA (Cayman Chemical) was performed according to manufacturer’s instructions. Briefly, plasma samples were hydrolysed to release any 8-isoprostane esterified in lipids. Plasma was then purified using 8-isoprostane Affinity Columns (Cayman Chemical, 401111). ELISA buffer, standards, samples, Tracer and Anti-serum were added to the plate and incubated for 18 h at 4°C. The plate was then washed, Ellman’s Reagent added and incubated for 2 h and read at 405 nm CLARIOstar plate reader (BMG Labtech).

### scRNA-seq dataset pre-processing

Gastrocnemius muscle from mice was collected and prepared for sequencing (n= 2/condition; one male, one female). Sexes were pooled for each condition. Library generation was performed using the 10X Genomics Chromium System (10X Genomics Inc San Fransico, CA) and reads aligned to mouse reference genome mm10-2020-A by Cell Ranger 6.0.2 to generate raw count matrices. Downstream analysis was performed in R 4.3.2. Quality control of datasets was performed independently using R packages Seurat 4.3.0^25^, SeuratObject 4.1.3 and Doublet Finder 2.0.3^26^ (Table S1). Datasets were integrated using Seurat method and normalisation of expression matrices was carried out using the “SCTransform” method.

### Sex determination of scRNAseq datasets

Sex was assigned to cells based on expression of Y-chromosome (*Ddx3y, Eif2s3y, Gm29650, Kdm5d, Uty*)^27^, X-chromosome genes (*Ddx3x, Eif2s3x, Kdm5c, Utx*) and X-chromosome silencing long-noncoding RNA *Xist*. As Y-chromosome genes are only expressed by male cells, but X-chromosome genes are expressed by both male and female cells, sex determination was carried out in two steps. Step one assigned a cell with a non-zero expression of *Xist* and zero expression of Y-chromosome genes as female, whereas a cell expressing one or more of the Y-chromosome genes with zero expression of *Xist* was assigned male. Step two used the expression of sex-assigned setting the parameters as follows: Y-gene cutoff as the lowest 10% of Y-chromosome gene expression in male cells. In female cells, Xist-cutoff is the median of *Xist* expression, lower X-gene cutoff is the lowest 20% of X-chromosome gene expression, and median X-gene cutoff is the median of X-gene expression. A cell with (a) non-zero expression of Xist and zero expression of Y-chromosome genes, and/or (b) sum of X-chromosome genes greater than the lower X-gene cutoff and zero expression of Y-chromosome genes, was assigned female. A cell with (a) expression of one or more of the Y-chromosome genes and zero expression of Xist, and/or (b) sum of Y-chromosome greater than Y-gene cutoff and Xist expression less than the Xist-cutoff, and/or (c) sum of Y-chromosome greater than Y-gene cutoff and X-gene expression less than the median X-gene cutoff, was assigned male. Cells that did not fit any of the above parameters were assigned as “Unknown” and filtered out of the dataset.

### Dataset integration, clustering and cluster identification

Datasets were normalized and scaled prior to integration using “IntegrateData” with an anchor set determined by “FindIntegrationAnchors” (method: canonical correlation analysis). Dimensionality of the data was captured by principal component analysis (PCA) within 45 dimensions, which were used for further analysis. Clustering was performed using Seurat’s “FindNeighbours” and “FindClusters” functions with a resolution of 0.5. Clusters were projected by uniform manifold approximation and projection (UMAP) by “RunUMAP”. Cluster identification was determined by “FindAllMarkers” function and by expression levels of known cell markers (Supplementary Figure 6A).

### Differential Gene Expression (DEG) Analysis and Gene Ontology

DEG analysis was conducted using Seurat’s “FindMarkers” function by the MAST test^28^. Genes were considered differentially expressed if they were expressed in at least 5% of cells tested and unadjusted p-value < 0.01. Sexually dimorphic genes were identified. Briefly, DEGs were identified between the ischemic and non-ischemic limbs of both male and female mice. Significant DEGs identified were then compared between ischemic limbs of male and female diabetic mice. Gene ontology analysis was conducted using the over-representation method in R package clusterProfiler 4.1.0^29^. Terms were considered significantly enriched adjusted p-value <0.05 and q-value <0.2.

### Tissue culture

Primary human male and female coronary artery ECs were purchased from American Type Culture Collection. Cells were cultured in ready-to-use MesoEndo Cell Growth Medium (Cell Applications, 212-500) or Endothelial Cell Growth Medium-2 BulletKit (Lonza, CC-3202) containing 5.5 mM D-Glucose (low glucose, LG). Human ECs were maintained at 37°C in a 5% CO_2_ humidified environment and not used beyond passage 8. Murine ECs were isolated from C57Bl6 mice and cultured as previously described^16^. Where required, media was supplemented with D-Glucose to a final concentration of 25 mM to mimic high glucose (HG) diabetic conditions *in vitro*.

### Proliferation assay

Male and female coronary artery ECs at equivalent passage were seeded into a 96-well plate at a density of 1x10^4^ cells/well in LG medium containing 5.5 mM D-glucose. The following day, the medium was changed into HG-containing medium and replaced every 2-3 days until 14 days later, where the cells were trypsinized and counted using the Beckman-Coulter-Z2 Particle-Count-and-Size-Analyser.

### Tubulogenesis assay

Pre-cooled 96-well plates were prepared by adding 100 μL of Growth Factor Reduced Matrigel® (Corning) to each well, followed by incubation at 37 °C for 30 minutes. Male and female coronary artery ECs at equivalent passage were trypsinized and seeded at a density of 1x10^4^ cells/Matrigel-coated well in 100 μL HG-containing medium. Images were acquired 6 h later using Olympus CKX41 microscope and ZEISS 105 colour camera at 4X magnification. The number of tubules and branch points were quantified using Wimasis analysis software (Spain).

### Seahorse Assay

Male and female coronary artery ECs at equivalent passage were seeded into 12-well plates at a density of 1.1x10^5^ cells per well. Cells were treated with HG media the following day, and media was changed every 2-3 days. On day 7, cells were trypsinized and seeded into a 0.1% gelatine-coated XF96 cell-culture-microplate (Agilent) at a density of 1x10^4^ cells per well and left for a further 7 days, with HG medium replaced every 2-3 days. Cells were washed with PBS prior to addition of XF RPMI media (180-μL; Agilent), supplemented with LG or HG, 1 mM pyruvate and 2 mM glutamine. Oxygen consumption rate (OCR) was determined using Seahorse-XF-Cell- MitoStress-Test kit (Agilent, California) and Seahorse-XFe96-analyser (Agilent), according to manufacturer’s instructions. Seahorse sensor cartridge was loaded with oligomycin (1.5 μM), carbonyl cyanide p-trifluoromethoxyphenylhydrazone (FCCP; 1 μM), rotenone (0.5 μM) and antimycin A (0.5 μM). Assays performed and analysed as described^30^.

### MitoSOX assay

Male and female coronary artery ECs at equivalent passage were seeded into a 96-well plate at a density of 1x10^4^ cells/well in LG medium. The following day, the medium was changed into LG or HG-containing medium and replaced every 2-3 days for 14 days. On day 13, cells were treated with either vehicle or 1 nM MitoQ for 24 hours. Following treatment, cells were washed with Hank’s Balanced Salt Solution (HBSS) and incubated with 5 μM MitoSOX red dye (Thermo Fisher Scientific) prepared in HBSS for 20 minutes at 37°C in the dark. Subsequently, cells were washed twice with HBSS to remove excess dye and imaged using a confocal microscope (excitation: 510 nm; emission: 580 nm; Zeiss 800). MitoSOX red fluorescence intensity was quantified using Image J software (version 1.53c, National Institute of Health, USA). Fluorescence intensity was normalized to cell number^31^.

### RNA isolation and quantitative polymerase chain reaction (qPCR)

RNA was isolated from mouse and human tissues using the RNeasy Fibrous Tissue Mini Kit (QIAGEN). For cells, male and female coronary artery ECs at equivalent passage were seeded into 12-well plates at a density of 1.1x10^5^ cells per well. Cells were treated with LG or HG media the following day, and the media was changed every 2-3 days. RNA was isolated 14 days later using TriReagent (Sigma-Aldrich). cDNA was generated using SensiFAST cDNA Synthesis Kit (Meridian Bioscience) and qPCR performed using iQ SYBR (Biorad, Australia) in a CFX38 thermocycler (Biorad, Australia). The ΔΔcT method normalized to β-actin or HPRT was used to assess mRNA changes. Primer sequences can be found in Supplementary Table S2.

### Statistics

Data are presented as mean ± SEM. Statistical analyses were performed using Student’s *t*-test, the Mann–Whitney *U* test, or one- or two-way ANOVA, followed, where applicable, by Bonferroni’s or Šídák’s post hoc tests. Unless stated otherwise, all analyses were carried out using GraphPad Prism version 10.01. A *P* value < 0.05 was considered statistically significant.

## RESULTS

### Female CLTI tissues have reduced arterial relaxation and microvessel numbers compared to male patients

To investigate sex differences in EC function(s) in diabetes-associated PAD, blood vessels and the surrounding tibialis anterior muscle were collected from diabetic CLTI patients following amputation (Figure 1A). Patient clinical characteristics are described in Table 1. A significant reduction in EC-dependent relaxation of isolated female vessels to acetylcholine was observed when compared to males, with no change in sodium nitroprusside-mediated relaxation (Figure 1B); a finding that was independent of vessel size (male 383.435 ±80.02 µm vs. female 269.9 ± 41.98 µm, *P*=0.196). Paraffin-embedded muscle tissues were stained and the number of microvessels <50 µm in diameter counted, normalized to myofiber number. Females had ∼50% fewer microvessels than male tissues, while myocyte numbers remained unchanged (Figure 1C-D). Female limb tissues also had a two-fold increase in mRNA expression of oxidative stress markers, NADPH oxidases *Nox-1* and *Nox-2* (Figure 1E). In contrast, plasma levels of 8-isoprostane, a biomarker that quantifies free radical-induced damage to lipids in circulation, were unchanged with sex (Figure 1F). These findings demonstrate altered EC responses within the local limb tissue environment of females with advanced diabetes-associated PAD.

**Figure 1.**
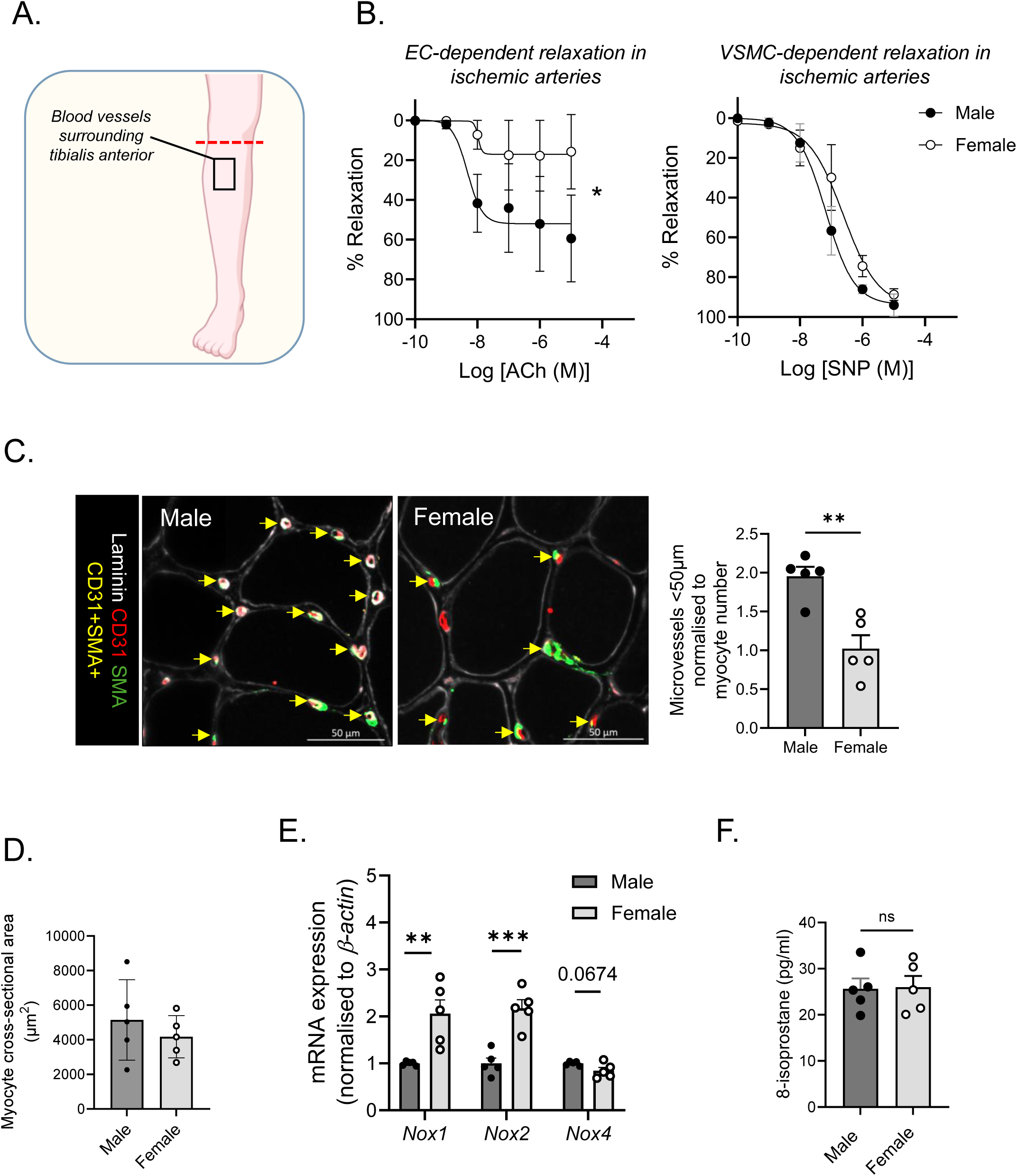
Amputated tissues from female patients with diabetes-associated PAD have reduced vasodilation and microvessel density. (A) Schematic illustrating site of tissue collection from below- and above knee amputations. (B) Myography showing increased endothelial dysfunction in female vs male patient arteries. *Left*, endothelial-dependent relaxation in response to acetylcholine (ACh). *Right*, No difference in vascular smooth muscle cell (VSMC), sodium nitroprusside (SNP)- mediated relaxation (n=3-5). (C) *Left,* representative image of microvessels in tibialis anterior (CD31^+^SMA^+^, yellow arrows). Laminin (myocytes, white), CD31 (ECs, red) and SMA (pericytes, green); scale bar 50 μm. *Right*, quantification (n=5/sex). (D) Myocyte area is unaltered with sex (n=5). (E) NADPH oxidase (*Nox*) mRNA marker expression in muscle measured by qPCR, normalized to β*-actin* (n=5). (F) 8-isoprostane levels in plasma remain unchanged with sex (n=5). Results are mean±SEM; two-way ANOVA, Student’s *t*-test or Mann–Whitney *U*-test; \**P*<0.05, \*\**P*<0.01, \*\*\**P*<0.001.

**Table 1.** Patient clinical characteristics.

| Characteristic | Male<br>n = 5 | Female<br>n = 5 |
| --- | --- | --- |
| Age (y) Mean (SD) | 67.6 (12.3) | 58.8 (8.8) |
| Body Mass Index Mean (SD) | 30.3 (5.3) | 32.4 (8.5) |
| Diabetic (%) | 5 (100) | 5 (100) |
| Hypertension (%) | 4 (80) | 2 (40) |
| Hypercholesterolemia (%) | 4 (80) | 3 (60) |
| Smoker (%) | 1 (20) | 4 (80) |
| HbaA1c (%) Mean (SD) | <sup>^</sup>9.40 (2.06) | 9.26 (4.00) |
| Footnote, <sup>^</sup>n=4. |  |  |

### Pre-clinical model of diabetes-associated PAD

To examine sex differences in preclinical diabetes-associated PAD, we established a new model in *Apoe^-/-^* atherosclerotic mice where diabetic parameters including plasma glucose and glucose tolerance were equivalent between the sexes (Figure S1; Table S3). As expected, mice weights were significantly lower with STZ treatment, and in female vs. male mice (Table S4). Female diabetic mice had ∼50% more plasma cholesterol, associating with increased liver weight (Table S3-4). No significant differences in atherosclerotic plaque size, media area or cap thickness of brachiocephalic arteries were observed with sex (Figure S2) however, aortic *Nox-2* mRNA was significantly elevated 4-fold in females, independent of diabetes (Figure S3A). *Icam-1*, *Il-1*β, and *Il-6* mRNA levels were also upregulated in female aortae ∼2-6-fold (Figure S3D-H).

### Female diabetes-associated PAD mice have reduced angiogenesis and arterial function, associating with elevated oxidative stress

To support the clinical significance of our findings, we measured EC functions in mice with diabetes-associated PAD. Vessels isolated from ischemic limbs of diabetic female mice showed a marked reduction in EC-dependent relaxation to acetylcholine compared to male vessels, along with mild impairment in SNP-mediated relaxation and unchanged plasma nitrate/nitrite levels (Figure 2A-D), indicating pronounced endothelial dysfunction with minor smooth muscle impairment, independent of systemic NO availability. Importantly, this impairment was not observed under non-diabetic non-ischaemic conditions (Figure 2C). Assessment of angiogenic capacity showed that diabetes impaired blood perfusion equally in both sexes (Figure 2E-G); however female diabetes-associated PAD mice had significantly fewer microvessel numbers in the gastrocnemius than male mice (Figure ); a finding also observed in female ECs *in vitro* (Figure 2I-J). Like human, ischemic skeletal muscle from diabetic female mice exhibited elevated mRNA expression of oxidative stress markers, *Nox-1* and *Nox-2* and markers of inflammation (*Icam-1*, *Vcam-1*, and *Il-6*) (Figure S4). Because diabetes impairs wound healing, we also assessed wound closure. Female diabetic *Apoe^-/-^*mice showed slower closure, reduced CD31 density and increased *Nox-2* mRNA (Figure S5). Together, these results reveal differences in EC responses and function in diabetes-associated PAD.

**Figure 2.**
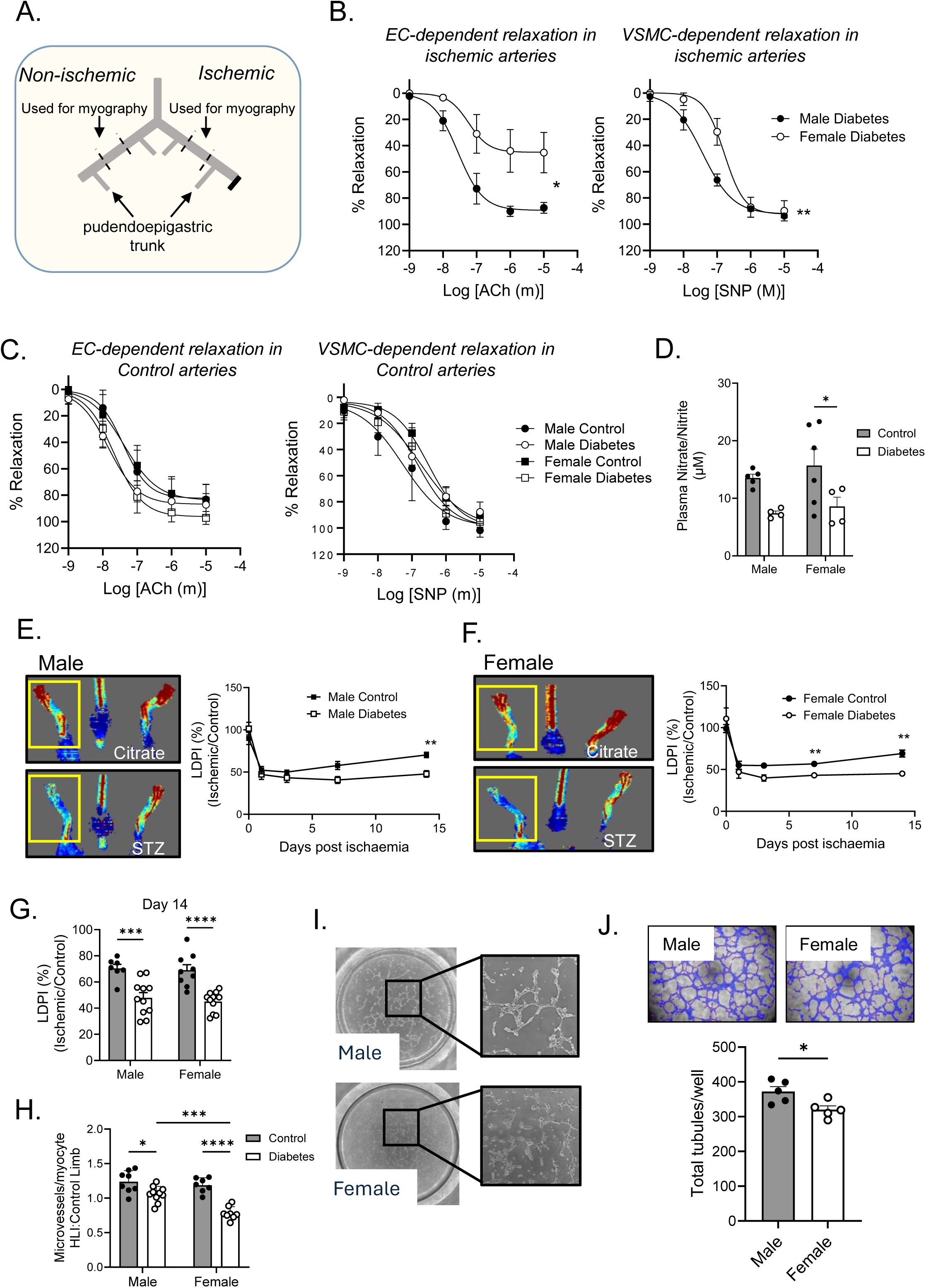
Diabetic female ischemic limbs have altered EC responses. (A) Schematic to show vessel collection for myography. (B) Myography showing increased endothelial dysfunction in female vs male arteries. *Left*, endothelial-dependent relaxation in response to acetylcholine (ACh). *Right*, sodium nitroprusside (SNP)-mediated relaxation (n=5-7). (C) Myography in non-injured vessels (N=5-7). (D) Plasma nitrite/nitrate levels (n=4-6). Laser Doppler imaging showing reduced blood perfusion over time with diabetes in (E) male and (F) female mice. *Left,* representative image of blood flow at 14 d. *Right,* quantification. (n=9-11). (G) Laser doppler perfusion index at 14 days (n=9-11). (H) Microvessel number (CD31+SMA+, <50 µm in diameter) in gastrocnemius tissues normalized to the number of myocytes and control non-ischemic limbs (n=9-11). (I) Tubule formation of male and female murine ECs shows an altered phenotype. (J) Tubulogenesis is reduced in female ECs under diabetic conditions. *Left,* Wimasis platform for vessel coverage and networks. *Right*, quantification (n=5/group). Results are mean±SEM; two-way ANOVA, Student’s *t*-test or Mann–Whitney *U*-test; \**P*<0.05, \*\**P*<0.01, \*\*\**P*<0.001 and \*\*\*\**P*<0.0001.

### ScRNA sequencing reveals sexual dimorphisms in ECs of diabetes-associated PAD mice

To examine the transcriptomic profiling of murine lower limbs after HLI, gastrocnemius was digested and cells filtered to isolate mononuclear cells, followed by scRNA-seq library preparation (Figure 3A). Unbiased clustering of 44,123 cells identified 24 cell populations including ECs, which were the second most abundant cell population (Figure 3B-C; Table S5). A closer analysis of this population revealed 3 subtypes (Figure 3D): EC1, represented by enriched expression for *Aqp1*, *Lpl* and *Adh1*, and marking mostly capillaries; EC2, represented by *Btnl9*, *Sema39* and *Rbp7*, marking mostly arteries and arterioles and EC3, represented by *Lrg1*, *Edh4* and *Ler3* expression and marking veins and venules (Figure E-F). Key genes were used to distinguish male and female ECs based on differential expression patterns (Figure 4A-B). Next, we identified sex-dimorphic genes differentially expressed in response to ischemia, with the most pronounced changes observed in the EC1 and EC2 populations (Figure 4C-D), which we prioritized for further analysis. GO term over- representation analysis revealed that female ECs showed upregulation of genes involved in mRNA metabolism, negative regulation of protein localization, and negative regulation of intracellular transport, while downregulated genes were linked to vasculogenesis, vascular processes in the circulatory system, and cell–substrate adhesion (Figure 4E). The top -up and -down regulated genes are shown in the volcano plot in Figure 4F and Tables S6-S7. Notably, mitochondrial complex I genes including *mt-Nd1*, *mt-Nd2*, *mt-Nd3*, *mt-Nd4*, *mt-Nd4l*, *mt-Nd5*, and *mt-Nd6*, which encode NADH dehydrogenase subunits, were markedly suppressed in ECs from female diabetes-associated PAD mice and in female amputation tissues (Figure 4G-H). Expression of *Cav1*—a gene recently linked to mitochondrial dynamics (ROS production, mitophagy, fission/fusion)^32–34^, and its adapter *Cavin2* were also reduced in female diabetic ischaemic ECs (Figure 4F, I; Table S7). Additional analyses revealed enriched terms related to mitochondrial function: regulation of mitochondrial membrane potential, regulation of mitochondrial organization, mitochondrial fission, mitochondrial localization, positive regulation of mitochondrial depolarization and positive regulation of establishment of protein localization to mitochondria.

**Figure 3.**
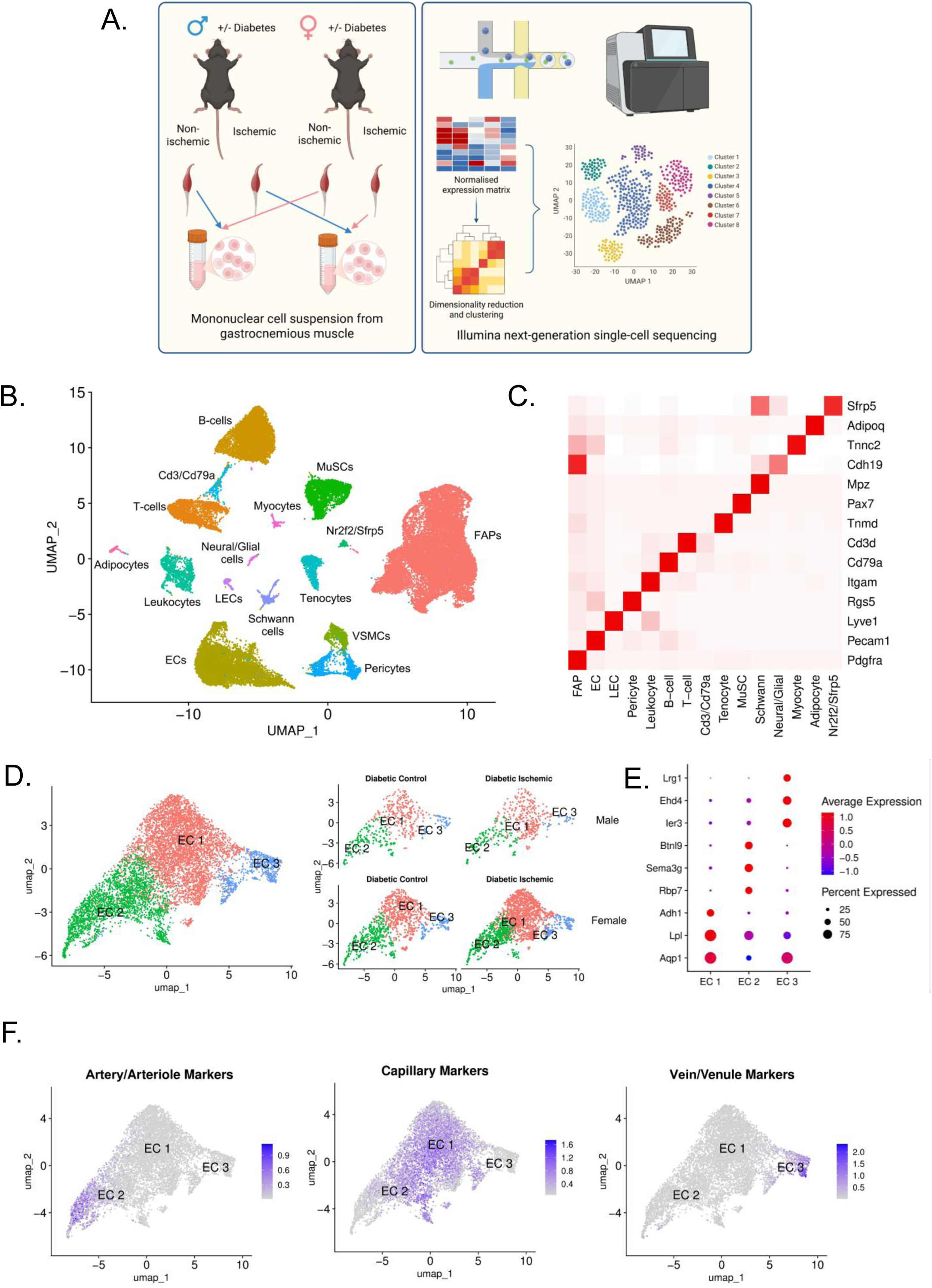
ScRNA-seq analysis of diabetes-associated PAD. (A) scRNA-seq workflow. (B) Principal component analysis followed by K-nearest neighbour clustering identified 24 populations including endothelial cells (ECs), lymphatic ECs (LECs), fibro-adipo progenitors (FAPs), vascular smooth muscle cell (VSMCs), pericytes, schwann cells, leukocytes, adipocytes, T cells, Cd/CD79a cells, B-cells, muscle stem cells (MuSCs), Nr2f2/Sfrp5 cells and neural/glial cells and myocytes. (C) Heat map of cell types using the most common cell markers. (D) scRNA-seq identified 3 EC clusters, which change with sex and diabetes. (E) Dot plot showing EC markers. (F) EC2 marks arterioles/arteries, EC3 marks mostly capillaries, while EC1 mark venules/veins.

**Figure 4.**
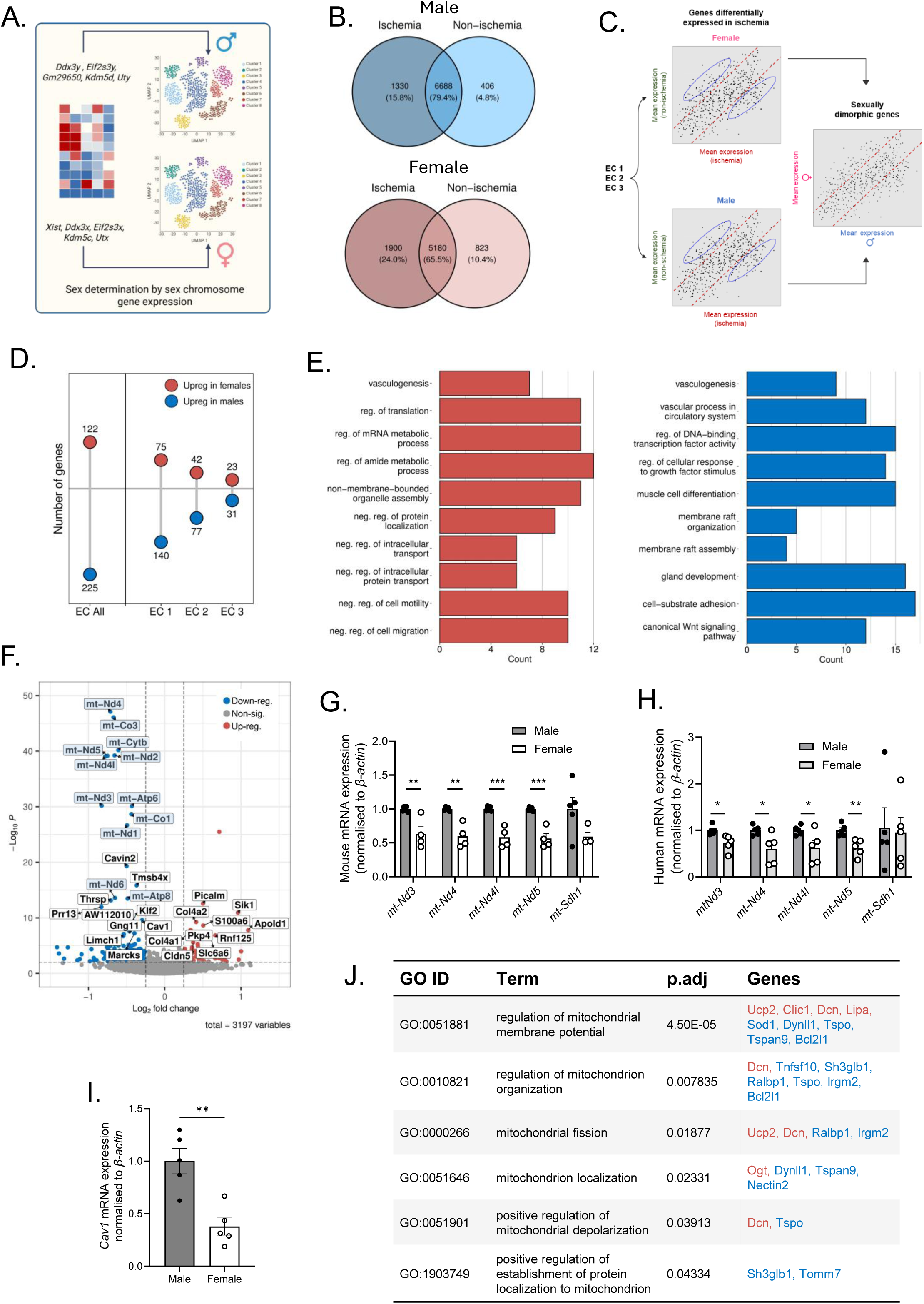
Sexual dimorphisms in ECs from diabetes-associated PAD. (A) Workflow. Sex chromosome expressions used to determine sexual dimorphisms in gene expression. (B) Venn diagram showing gene changes in in male (*upper*) and female (*lower*) diabetic ischemic vs non-ischemic gastrocnemius. (C) Workflow to show sexually dimorphic genes and the ischemic response in diabetes. (D) Lollipop plot depicting the number of genes upregulated in male and female diabetic ischemic limbs. (E) Over-representation analysis showing positive (*left*) and negative (*right*) enriched GO terms in female ischemic diabetic limbs. (F) Volcano plot showing the top up and down-regulated gene changes. Light blue boxes highlight mitochondrial genes suppressed in female diabetes-associated PAD. Multiple complex I mitochondrial genes are suppressed in diabetic ischemic limbs of (G) female mice and (H) women undergoing amputation. (I) *Cav1* mRNA expression is reduced in female vs male amputated skeletal muscle; mRNA normalized to β-actin (n=5). (J) Further analyses identifying additional GO terms related to mitochondria, and the genes that are altered. Red, up-regulated; blue, down-regulated.

### Mitochondrial function is reduced in diabetic female ECs

Various strategies were employed to evaluate sex-dependent EC mitochondrial function(s) under diabetic conditions *in vitro*. Real-time Seahorse bioenergetic analysis revealed that female ECs exposed to HG, but not LG for 14 days had a reduced oxygen consumption rate (Figure 5A), and supporting this, Electron Transport Chain Complex I genes, *Nd3*, *Nd4*, *Nd4l*, and *Nd5* were more robustly reduced in female than male ECs under diabetic conditions (Figure 5B). To see if a decrease in respiration under HG conditions reflected fewer mitochondria, we measured mitochondrial mass. Under basal conditions female ECs have been shown to have more mitochondria^35^. We too observed greater mitochondrial content in female vs male ECs; a response no longer evident under diabetic conditions (Figure 5C). MitoSOX staining, indicative of mitochondrial ROS, increased in both male and female ECs, but the rise was markedly greater in female ECs under diabetic conditions (Figure 5D). Because our scRNA-seq data revealed enriched terms related to mitochondrial permeability transition pore, organization, and localization, along with altered expression of associated genes (Figure 4G), we validated expression of *Tspan9* and *Dynll1*. TSPAN9, a tetraspanin, regulates vesicle trafficking and mitochondrial interactions, while DYNLL1 controls mitochondrial distribution, dynamics, and function via dynein-mediated transport; both were reduced in female diabetic PAD limbs (Figure 5E). Collectively, these indicate sex-dependent differences in EC mitochondrial function with diabetes.

**Figure 5.**
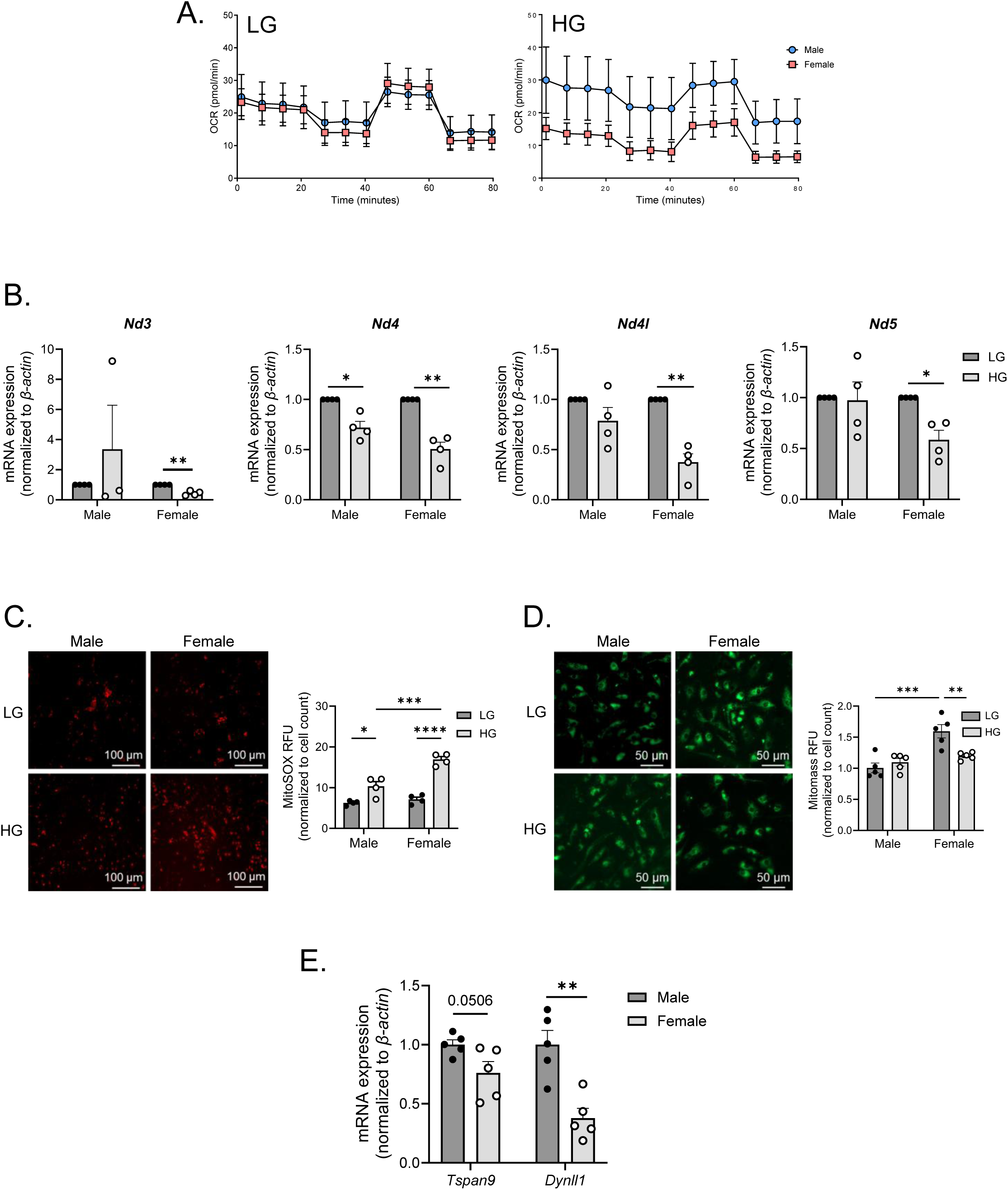
Mitochondrial function is reduced in female ECs under diabetic conditions. (A) Oxygen consumption rate between male and female ECs under (*left*) low glucose (LG), and (*right*) high glucose (HG)-containing conditions for 14 days (n=4). (B) mRNA expression of complex I genes are reduced in female ECs under 14-day LG and HG conditions. Gene expression as normalized to β*-actin* (n=4). (C) Mitotracker green showing mitochondrial abundance between sex under LG and HG-containing conditions. *Left*, representative image. *Right*, quantification (n=5). Scalebar represents 100 µm. (D) MitoSOX staining is increased with diabetes (14 days) and more so in female ECs. *Left*, representative image. *Right*, quantification (n=4). Scalebar represents 100 µm. (E) mRNA expression of *Tspan9* and *Dynll1*, normalized to β-actin (n=5). Soleus muscle of diabetes-associated PAD mice. Results are mean±SEM; two-way ANOVA, Student’s *t*-test or Mann–Whitney *U*-test; \**P*<0.05, \*\**P*<0.01, \*\*\**P*<0.001 and \*\*\*\**P*<0.0001.

### MitoQ restores EC function(s) in diabetes-associated PAD

MitoQ is a mitochondrial-targeted antioxidant designed to selectively reduce oxidative damage within the mitochondria. We investigated whether MitoQ could impact diabetes-associated PAD in a sex-specific manner. To assess this, a shorter diabetes-associated PAD model was used where MitoQ was delivered via osmotic pumps at 10 µg/day for 2 weeks post-HLI (Figure 6A). MitoQ significantly improved EC-dependent relaxation to acetylcholine in female, but not male arteries, with no significant effect on SNP-mediated relaxation (Figure 6B-D). MitoQ intervention also increased microvessel density in gastrocnemius muscle of female mice with no change in blood perfusion (Figure 6E-F). *In vitro*, MitoQ reduced EC mitochondrial superoxide production in both sexes, with more profound effects in female ECs exposed to 14-day HG conditions (Figure 6G) but had no significant effect on mitochondrial content (Figure 6H). These findings suggest that MitoQ supplementation improves mitochondrial dysfunction, particularly in female diabetes-associated PAD, restoring EC function(s).

**Figure 6.**
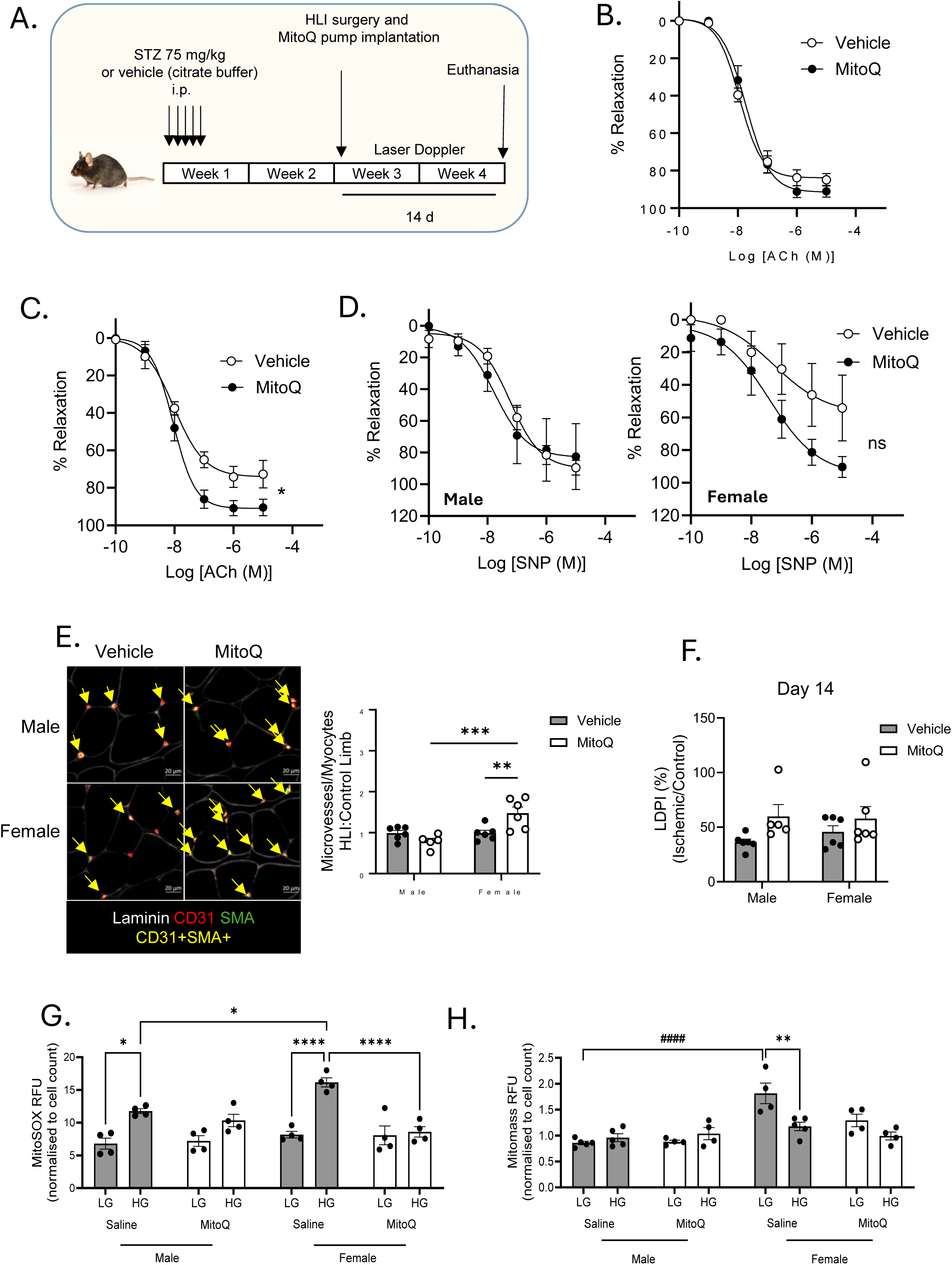
MitoQ restores EC function in diabetes-associated PAD. (A) Schematic showing a shorter diabetes-associated PAD model with MitoQ (10 µg/day) or vehicle supplementations with Alzet pumps for 14 days after hindlimb ischemia (HLI). MitoQ has no effect on (B) male diabetic ischemic vessels (n=5-6) but improves EC-dependent vasorelaxation in (C) female diabetic ischemic vessels (n=6). (D) There is no change with SNP-mediated relaxation (n=5-6). (E) MitoQ increases the number of microvessels (CD31+SMA+, <50 µm in diameter) in female ischemic limbs (n=6), (F) with no change to Laser doppler perfusion index at 14 days (n=5-6). (G) MitoQ (1 nM) has no effect on mitochondrial abundance in 14-day LG and HG-containing conditions (n=4). (H) MitoSOX staining is reduced with MitoQ administration (1 nm) under HG-containing conditions (14 days) (n=4). Results are mean±SEM; two-way ANOVA; \**P*<0.05, \*\**P*<0.01, \*\*\**P*<0.001 and \*\*\*\**P*<0.0001.

## DISCUSSION

The 2025 International Diabetes Federation Diabetes Atlas reports that ∼11% or ∼589M of the adult population (20-79 years) is living with diabetes world-wide; a number predicted to rise to 853M in the next 25 years^36^. Diabetes is a key driver of PAD. Integrating sex as a biological variable is crucial to advancing equitable and effective care - especially as the burden of PAD rises in ageing, and diabetic female populations, who experience worse outcomes to treatment^37^. In this study, we discovered an exciting novel mechanism implicating mitochondrial endothelial dysfunction as a driving force for poorer vascular health in females with diabetes-associated PAD. Specifically, five critical, sex-specific alterations were identified: (i) vasodilatory responses were significantly blunted in vessels isolated from females - both in mice and humans with diabetes - compared to their male counterparts; (ii) angiogenic capacity was reduced in ischemic limbs of diabetic female mice, with reduced microvessel density also evident in limb tissues from female patients undergoing amputation; (iii) localised oxidative stress markers were significantly elevated in females, (iv) with female murine ECs exhibiting distinct gene signatures highlighting disrupted mitochondrial dynamics, supported by reductions in mitochondrial activity and increased mitochondrial ROS, driving endothelial dysfunction in diabetes. Remarkably, (v) treatment with the mitochondria-targeted antioxidant MitoQ enhanced vasodilation, restored the endothelium’s angiogenic potential, and reduced oxidative stress in female diabetes-associated PAD.

ECs are key regulators of vascular tone and angiogenesis. Strikingly, female vessels from diabetes-associated PAD mice and humans exhibited impaired vasodilation and lower microvessel density compared to males. Premenopausal women generally have greater endothelial function compared to men of a similar age^38^, largely driven by estrogen, which upregulates endothelial nitric oxide synthase, increases NO bioavailability, and improves vascular smooth muscle sensitivity to NO. Women also tend to have less oxidative stress and more favourable lipid profiles before menopause^39,40^, which protects endothelial function. After menopause, the loss of estrogen is linked to a marked reduction in vasodilatory capacity^41^ and falls to similar levels to those seen in age-matched men^42^. Similarly, angiogenic ability is reduced in women post-menopause^43^. The average age of women in our PAD cohort (∼59 years) implies that postmenopausal estrogen decline could contribute to reduced vascular tone and microvessel density; however, young adult female mice with diabetes-associated PAD representing young to middle age, exhibited a comparable phenotype. Likewise, *in vitro* studies of ECs cultured in hormone-free media demonstrated persistent functional deficits in female cells, supporting a hormone-independent mechanism; findings that require further investigation.

Interestingly, sex differences in capillary density in PAD remain inconsistent. For example, no sex-based differences were observed among PAD patients who were asymptomatic or had intermittent claudication^44^, including those with diabetes (personal communication, Mary McDermott). In contrast, another study found that women had significantly lower capillary density when measured as the EC-to-fibre ratio, but not when assessed by capillaries per tissue area^45^. In our study, we quantified microvessels <50 µm in diameter^46^, normalized to myocyte number. We find significant reductions in these vessels in female patients. Supporting our results, a Matrigel plug assay revealed significantly fewer CD31[ cells in female versus male plugs following FGF-2–induced angiogenesis^47^, and likewise, in a preclinical PAD model, female ischemic limbs exhibited blunted angiogenic responses to HLI^47^. However, the majority of studies assessing capillary or microvascular density in advanced PAD, such as CLTI, have not incorporated sex-based tissue comparisons^48,49^, limiting ability to detect sex-specific vascular changes. Methodological differences in assessing microvessels are likely to contribute to the variability in reported findings. In our system, we observed fewer microvessels in skeletal muscle from female diabetes-associated PAD patients, from mice and ECs *in vitro*.

Mitochondrial dysfunction is evident in PAD^50,51^. Reduced ATP availability disrupts cellular energy balance, fuels inflammation and oxidative stress, triggers cell death, and hampers both wound repair and new blood vessel formation; factors that collectively undermine tissue function and contribute to the clinical decline. Indeed, reduced mitochondrial DNA and content are associated with cardiovascular disease and predict mortality and adverse events in PAD patients^52^. OMICs analyses of gastrocnemius muscle by others revealed transcriptomic and proteomic signatures of mitochondrial damage^53^, while functional studies demonstrated impaired respiration in both patients and murine ischemic muscle, accompanied by elevated oxidative stress^51,54,55^. Interestingly, direct evidence linking both endothelial and mitochondrial dysfunction in limb tissues in PAD exist. Isolated arterioles from PAD patients exhibited reduced endothelium-dependent vasodilation, correlating with impaired mitochondrial function in tissues^50^. However, to our knowledge sex differences in EC mitochondrial function(s), particularly in diabetes have not been described. Our transcriptomic analysis revealed that ECs from male and female diabetes-associated PAD mice have markedly different gene expression programs; with female ECs showing pronounced enrichment in mitochondrial pathways, and marked suppression of mitochondrial complex genes, suggesting sex differences in mitochondrial function. Indeed, we observed reduced mitochondrial content, respiration, and increased oxidative stress in female vs male ECs under chronic diabetic conditions. Several genes of interest were identified from our scRNA-seq analysis, including Cav1: critical for caveolae formation. Cav1 has pleiotropic effects. For example, *Cav1^-/-^* mice develop cardiomyopathy and pulmonary hypertension with elevated NO^56^. Important in this setting Cav1 was shown to regulate eNOS activation, barrier function, and angiogenesis^57^, as well as mitochondrial respiration, dynamics, calcium signalling, and mitophagy^32,33^. In ECs, loss or dysregulation of Cav1 impaired mitochondrial membrane potential, increased ROS, and disrupted fission/fusion balance and calcium homeostasis, contributing to endothelial dysfunction^34^. However, Cav1’s role in diabetes-associated PAD is unknown. We found reduced *Cav1* mRNA in ECs and ischaemic limb tissues of diabetic female mice, as well as in female patients. The role of Cav1 suppression as a potential mediator of mitochondrial dysfunction in female diabetes-associated PAD is yet to be defined. Interestingly, Cav1 knockdown in cancer cells increased oxidative phosphorylation^32^, and female EC in an obesogenic study were enriched for oxidative phosphorylation^58^, suggesting that metabolic alterations may represent a common, sex-specific mechanism influencing cell fate. Our findings indicate that enhanced mitochondrial dysfunction in female ECs following injury and diabetes may underlie the heightened vascular vulnerability observed in women with PAD.

MitoQ is a mitochondria-targeted antioxidant in which a modified coenzyme Q10 is linked to a lipophilic triphenylphosphonium cation through a 10-carbon alkyl chain, allowing it to preferentially accumulate within mitochondria. MitoQ was reported to attenuate age-related arterial dysfunction in mice^59,60^, and more recently administered to healthy older individuals, greater than 60 years of age^60^. Chronic MitoQ supplementations for 6 weeks in these individuals, improved flow mediated dilatation, as a measure of endothelial function, when compared to placebo^60^. MitoQ also reduced mitochondrial ROS levels and increased NO bioavailability^60^. Related to this study, acute MitoQ administration boosted flow-mediated dilation and walking performance in PAD patients, though sex-specific effects were not stratified^61^. Although MitoQ’s sex-dependent effects are not fully defined, a preclinical study showed it increased oxygenation more effectively in female fetal placentae under hypoxia^62^, hinting at greater benefits in females. Indeed, at a single concentration of 10 µg/day for 14 days, we observed a marked enhancement of EC function, namely, vasodilation and angiogenic capacity in female vs male diabetes-associated PAD mice. Female ECs under diabetic conditions also exhibited reduced mitochondrial oxidative stress. These findings suggest that targeting mitochondrial dysfunction with MitoQ could be a promising strategy to restore vascular health in PAD, with potential sex-specific therapeutic implications, however further studies are needed to identify sex-specific effects and optimal dosing.

Our findings provide significant insights into sex differences and EC function(s) in diabetes-associated PAD however several limitations should be acknowledged. Most notably, our cohort included only patients with advanced disease, so it remains unclear whether these sex-related differences are also present in asymptomatic individuals or those with intermittent claudication. Second, the sample size was small, with a maximum of five patients per sex. Third, the HLI model has inherent constraints, representing an acute injury that induces severe ischemia rather than chronic atherosclerotic disease, and *Apoe^-/-^*mice do not develop atherosclerosis in limb vessels under these conditions. Finally, we did not evaluate the potential contribution of sex chromosomes to the observed differences, nor the influence of maternally inherited mitochondrial DNA on mitochondrial function. Thirteen mitochondrial DNA-encoded proteins including Complex I (ND1, ND2, ND3, ND4, ND4L, ND5, ND6), Complex III (Cytochrome B), Complex IV (COX1-3) and Complex V (ATP6, ATP8) are transmitted exclusively from the mother. Because our studies were performed in siblings from both sexes, the observed changes are unlikely to reflect maternal inheritance and most mitochondrial proteins (>1000) are encoded by nuclear DNA, supporting a nuclear driven mechanism. Nevertheless, our findings highlight mitochondrial dysfunction as a key contributor to endothelial impairment in diabetic females, suggesting intrinsic, sex-specific vulnerabilities in mitochondrial regulation. Recognising and addressing these differences is crucial for improving our understanding of female-predominant vascular risk and for guiding the development of targeted therapeutic strategies.

## ACKNOWLEDGMENTS

We would like to acknowledge Ms Brooke Waldram and Mr Millard Phitphatphol for their technical expertise.

