## supplementary Tables and Figs for "Mitochondrial Dysfunction in Endothelial Cells Drives Greater Vascular Impairment in Females with Diabetes-Associated Peripheral Artery Disease"

#### SUPPLEMENTARY TABLES AND FIGURES

#### SUPPLEMENTARY TABLES

##### Table S1. Data pre-processing steps for scRNA-seq.

| Sample | Quality Control and doublet discrimination | nPcs and resolution for clustering | Expected % doublets According to 10X Recommendation | Doublet Finder Results |
|---|---|---|---|---|
| Non-diabetic Control | Started with 18960 cells<br>1. nFeature_RNA > 200 & nFeature_RNA < 5000 & percent.mt < 25<br>18186<br>2. percent.mt < 15<br>17246<br>3. Remove cluster 4<br>15913<br>4. Remove cluster 8<br>15354 | Dims = 1:20<br>Res = 0.5 | 0.8*15.354~12.2% | Singlet = 14306<br><br>Doublet = 1048 |
| Non-diabetic Ischemic | Started with 13766 cells<br>1. nFeature_RNA > 200 & nFeature_RNA < 5000 & percent.mt < 25<br>13498<br>2. percent.mt< 20<br>13387 | Dims = 1:20<br>Res = 0.5 | 0.8*13.387~10.71% | Singlet = 12469<br><br>Doublet = 918 |
| Diabetic Control | Started with 15436 cells<br>1. nFeature_RNA > 200 & nFeature_RNA < 6000 & percent.mt < 25<br>14701<br>2. percent.mt < 15<br>14222<br>3. Remove cluster 5<br>13369 | Dims = 1:20<br>Res = 0.5 | 0.8*13.369~10.70% | Singlet = 12473<br><br>Doublet = 896 |
| Diabetic Ischemic | Started with 18084 cells<br>1. nFeature_RNA > 200 & nFeature_RNA < 6000 & percent.mt < 25<br>17842<br>2. percent.mt < 15<br>17497<br>3. Remove cluster 7<br>16793 | Dims = 1:20<br>Res = 0.5 | 0.8*16.793~13.43% | Singlet= 15868<br><br>Doublet = 1107 |

### Table S2. qPCR primer sequences

| Human | | |
|---|---|---|
| | **Forward 5'-3'** | **Reverse 5'-3'** |
| *Icam-1* | GTATGAACTGAGCAATGTGCAAG | GTTCCACCCGTTCTGGAGTC |
| *Vcam-1* | TTTGACAGGCTGGAGATAGACT | TCAATGTGTAATTTAGCTCGGCA |
| *Il-1b* | AAACAGATGAAGTGCTCCTTCC | AAGATGAAGGGAAAGAAGGTGC |
| *Il-18* | TCTTCATTGACCAAGGAAATCGG | TCCGGGGTGCATTATCTCTAC |
| *Il-6* | ACTCACCTCTTCAGAACGAATTG | CCATCTTTGGAAGGTTCAGGTTG |
| *Tnf-α* | GTTGTAGCAAACCCTCAAGCTG | GAGGTACAGGCCCTCTGATG |
| *Mcp-1* | CAGCCAGATGCAATCAATGCC | TGGAATCCTGAACCCACTTCT |
| *Nox1* | TTCACCAATTCCCAGGATTGAAGTGGATGGTC | GACCTGTCACGATGTCAGTGGCCTTGTCAA |
| *Nox2* | GTCACACCCTTCGCATCCATTCTCAAGTCAGT | CTGAGACTCATCCCAGCCAGTCAGGTAG |
| *Nox4* | CTGGAGGAGCTGGCTCGCCAACGAAG | GTGATCATGAGGAATAGCACCACCACCATGCAG |
| *mt-Nd3* | CCACAACTCAACGGCTACAT | TGGGTGTTGAGGGTTATGAG |
| *mt-Nd4* | CCCTCGTAGTAACAGCCATTCTC | CGACTGTGAGTGCGTTCGTAGT |
| *mt-Nd4l* | CCCTCGTAGTAACAGCCATTCTC | CGACTGTGAGTGCGTTCGTAGT |
| *mt-Nd5* | GCCTAGCATTAGCAGGAATA | GAGTTTTAGGTAGAGGGGGA |
| *mt-Sdh1* | GAGATGTGGTGTCTCGGTCCAT | GCTGTCTCTGAAATGCCAGGCA |
| *b-actin* | AGCCATGTACGTAGCCATCC | CTCTCAGCTGTGGTGGTGAA |
| *18S* | CGGCTACCACATCCAAGGAA | GCTGGAATTACCGCGGCT |
| **Mouse** | | |
| *Icam-1* | GTGATGCTCAGGTATCCATCCA | CACAGTTCTCAAAGCACAGCG |
| *Vcam-1* | TTGGGAGCCTCAACGGTACT | GCAATCGTTTTGTATTCAGGGGA |
| *Il-1b* | GTTTCTGCTTTCACCACTCCA | GAGTCCAATTTACTCCAGGTCAG |
| *Ill-18* | GACTCTTGCGTCAACTTCAAGG | CAGGCTGTCTTTTGTCAACGA |
| *Il-6* | CTGCAAGAGACTTCCATCCAG | AGTGGTATAGACAGGTCTGTTGG |
| *Tnf-α* | CATCTTCTCAAAATTCGAGTGACAA | TGGGAGTAGACAAGGTACAACCC |
| *Mcp-1* | GCTGGAGCATCCACGTGTT | ATCTTGCTGGTGAATGAGTAGCA |
| *Nox1* | TCTCCAGCCTATCTCATCCTGA | GCTGCATCCATCACTGTCATGTT |
| *Nox2* | AGCTATGAGGTGGTGATGTTAGTGG | CACAATATTTGACCAGACAGACTTGAG |
| *Nox4* | CCCAAGTTCCAAGCTCATTTCC | TGGTGACAGGTTTGTTGCTCCT |
| *mt-Nd3* | TAGTTGCATTCTGACTCCCCCA | GAGAATGGTAGACGTGCAGAGC |
| *mt-Nd4* | CGCCTACTCCTCAGTTAGCCA | TGATGTGAGGCCATGTGCGA |
| *mt-Nd4l* | AGCTCCATACCAATCCCCATCAC | GGACGTAATCTGTTCCGTACGTGT |
| *mt-Nd5* | GGCCCTACACCAGTTTCAGC | AGGGCTCCGAGGCAAAGTAT |
| *mt-Sdh1* | GAGATACGCACCTGTTGCCAAG | GGTAGACGTGATCTTTCTCAGGG |
| *β-actin* | AGCCATGTACGTAGCCATCC | CTCTCAGCTGTGGTGGTGAA |

**Table S3. Diabetic mice plasma chemistries**

| | Male | | Female | |
|---|---|---|---|---|
| | Control | Diabetes | Control | Diabetes |
| Glucose (mmol/L) | 6.19 ± 0.37 | 18.82 ± 1.71*** | 5.67 ± 0.42 | 17.46 ± 2.56**** |
| Cholesterol (mmol/L) | 7.01 ± 0.371 | 15.56 ± 1.78 | 5.64 ± 0.71 | 35.15 ± 9.23*** # |

**Table S4. Diabetic mice body and organ weights**

| | Male | | Female | |
|---|---|---|---|---|
| | Control | Diabetes | Control | Diabetes |
| Body weight (g) | 30.44 ± 0.53 | 20.91 ± 0.61**** | 24.62 ± 0.86#### | 18.28 ± 0.64**** # |
| Liver per body weight (mg/g) | 39.30 ± 0.41 | 52.69 ± 1.30** | 42.27 ± 1.87 | 65.50 ± 3.08**** ### |
| Spleen per body weight (mg/g) | 3.00 ± 0.12 | 2.82 ± 0.24 | 3.78 ± 0.22 | 6.83 ± 1.47 ## |
| Kidney per body weight (mg/g) | 6.21 ± 0.23 | 10.01 ± 0.43**** | 6.34 ± 0.30 | 8.96 ± 0.63** |
| Retro fat per body weight (mg/g) | 2.60 ± 0.72 | 0.00 ± 0.00 | 6.94 ± 1.57## | 0.63 ± 0.29**** |
| Fat per body weight (mg/g) | 7.69 ± 1.48 | 0.00 ± 0.00**** | 9.71 ± 1.35 | 0.85 ± 0.67**** |

Footnote. * Difference within sex; # difference between sex; 2way ANOVA.

**Table S5. Known cell markers for scRNA-seq cluster identification.**

| Cluster Cell Type | Markers |
|---|---|
| B-cell | *Ms4a1*<br>*Cd19* |
| Endothelial cell | *Pecam1*<br>*Cdh5*<br>*Emcn* |
| Fibro-adipose progenitor | *Pdgfra*<br>*Col1a1*<br>*Apod*<br>*Lum*<br>*Fn1* |
| Leukocyte | *Adgre1*<br>*Fcgr1*<br>*Csf1r*<br>*Itgam* |
| Lymphatic endothelial cell | *Lyve1* |
| Pericyte | *Vtn*<br>*Pdgfrb*<br>*Rgs5* |
| Muscle Stem Cell | *Pax7*<br>*Myf5* |
| Schwann cell | *Mbp*<br>*Plp1* |
| Myocyte | *Tnnc2* |
| T cell | *Cd3c*<br>*Cd3g* |
| Tenocyte | *Tnmd* |
| Vascular Smooth Muscle Cell | *Acta2*<br>*Myh11* |

**Table S6. Top 10 sexually dimorphic genes upregulated in female diabetes-associated PAD mice.**

| Gene | Fold Change | p-value | Function | Reference |
|---|---|---|---|---|
| Picalm | 1.416283 | 2.56E-13 | Involved in clathrin coated pit internalization of membrane proteins; Regulates Aβ BBB transcytosis and clearance in brain endothelial cells | 1-3 |
| Sik1 | 1.944882 | 1.21E-11 | Upregulation protects again brain microvascular endothelial cells against oxygen-glucose deprivation, reoxygenation-induced injury; co-upregulation in cancer inhibits angiogenesis | 4,5 |
| Col4a2 | 1.33022 | 5.62E-10 | Structural component of basement membrane; associations between angiogenesis in skeletal muscle after HIIT | 6 |
| S100a6 | 1.418602 | 2.33E-09 | Regulates endothelial cell progression through the cell cycle; protects the endothelial barrier against calcium entry-induced disruption | 7,8 |
| Col4a1 | 1.264961 | 1.58E-08 | Structural component of basement membrane; associations between angiogenesis in skeletal muscle after HIIT | 6 |
| Apold1 | 2.13509 | 1.66E-08 | Localized APOLD1 to EC contacts and to Weibel-Palade bodies; associates with von Willebrand factor tubules. Role in maintaining cell junction-cytoskeletal interface, endothelial cell permeability; role in pathological angiogenesis | 9,10 |
| Pkp4 | 1.457181 | 9.35E-08 | Cell adhesion and cytoskeletal organisation; target gene of a proangiogenic miRNA | 11,12 |
| Rnf125 | 1.583407 | 1.43E-07 | Promotes p53 degradation; | 13 |
| Slc6a6 | 1.513552 | 1.63E-07 | Gated taurine transporter; uptake of gamma-aminobutyric acid (GABA); prevents VSMC Proliferation and Migration via Wnt, β-Catenin signaling | 14,15 |
| Cldn5 | 1.294802 | 2.64E-06 | Integral membrane protein; tight junctions; expression regulates EC permeability | 16,17 |

**Table S7. Top 10 sexually dimorphic genes downregulated in female diabetes-associated PAD mice.**

| Gene | Fold Change | p-value | Function | Reference |
|---|---|---|---|---|
| Cavin2 | -1.41477 | 4.81E-20 | Regulates the activity and stability of endothelial nitric-oxide synthase (eNOS) in angiogenesis | 18 |
| Tmsb4x | -1.29106 | 1.40E-16 | Promotes endothelial differentiation potential of human adipose-derived stem cells by up-regulating various angiogenic genes | 19 |
| Thrsp | -1.63318 | 7.44E-14 | Role in mitochondrial maintenance and sphingolipid metabolism in adipocytes | 20 |
| Prr13 | -1.77971 | 1.09E-12 | Downregulates thrombospondin 1 which prevents angiogenesis by inducing endothelial cell death | 21 |
| Cav1 | -1.22786 | 2.54E-10 | Binding with eNOS controls expression and function; role in endothelial permeability and thrombo-inflammation | 22,23 |
| Klf2 | -1.33876 | 3.28E-10 | Zinc-finger transcription factor; provides protection after endothelial cell injury; pathway for regulating eNOS | 24,25 |
| AW112010 | -1.5174 | 1.73E-09 | Promotes mitochondrial biogenesis; regulate expression of IL-10 | 26,27 |
| Gng11 | -1.37738 | 2.76E-08 | Promotes senescence pathways | 28,29 |
| Marcks | -1.31675 | 6.07E-08 | Regulates EC proliferation; role in insulin-dependent endothelial signaling to PIP(2); actin assembly and directed cell movement | 30,31 |
| Limch1 | -1.4515 | 8.67E-08 | Regulates and suppresses cell migration through regulating non-muscle myosin | 32 |

**SUPPLEMENTARY FIGURE LEGENDS**

**Figure S1. Preclinical model of diabetes-associated PAD.** (A) Schematic depicting the model. STZ, streptozotocin; GTT, glucose tolerance test; HLI, hindlimb ischemia. (B) Weekly unfasted plasma glucose levels (n=8-12). (C) Glucose tolerance tests in diabetes-assoiated PAD mice (n=8-12). Results are mean±SEM; two-way ANOVA; ****$P$<0.0001.

**Figure S2. Plaque size in unchanged with sex.** (A) Representative image of hematoxylin and eosin staining of brachiocephalic arteries. Scale bar = 20 μm. (Quantification of (B) plaque and (C) media area, normalized to artery area (n=7-8). Quantification of necrotic core area normalized to plaque area (n=6-8). Results are mean±SEM; two-way ANOVA or Mann–Whitney *U*-test; *$P$<0.05 and ***$P$<0.001.

**Figure S3. Female mice have increased expression of aortic oxidative stress and inflammatory markers.** qPCR analysis of (A-C) *Nox2*, *Nox1*, *Nox4;* (D-E) adhesion molecules *Icam-1* and *Vcam1*; inflammasome markers (E-H) *Il-1β*, *Il-18* and *Il-6*; and general inflammatory cytokine (I) *Tnf-α* and (J) *Mcp-1 mRNA expression.* Normalized to β-actin (n=5-7). Results are mean±SEM; two-way ANOVA; *$P$<0.05, **$P$<0.01, ***$P$<0.001 and ****$P$<0.0001.

**Figure S4. Female ischemic limbs have increased oxidative stress and inflammatory gene changes.** qPCR analysis of (A-C) *Nox2*, *Nox1*, *Nox4;* (D-E) adhesion molecules *Icam-1* and *Vcam1*; inflammasome markers (E-H) *Il-1β*, *Il-18* and *Il-6*; and general inflammatory cytokine (I) *Tnf-α* and (J) *Mcp-1 mRNA expression.* Normalized to β-actin (n=4-7). Results are mean±SEM; two-way ANOVA; *$P$<0.05, **$P$<0.01 and ***$P$<0.001.

**Figure S5. Angiogenesis and wound healing are reduced in female diabetic mice.** (A) Schematic depicting the wound healing model. (B) Blood flow is evident in female wounds. *Left*, representative doppler images of wounds. Red depicting more blood flow; blue representative of less blood flow. *Right*, Laser doppler perfusion index (LDPI) over time (n=10). (C) CD31 florescence intensity as a measure of capillary density, normalized to area (n=10). (D) mRNA expression of *Nox1*, *Nox2* and *Nox4* in wounds, normalized to β-actin (n=7). Results are mean±SEM; two-way ANOVA or Students' *t*-test ***P*<0.01.

**Figure S1: Preclinical model of diabetes-associated PAD**

A.

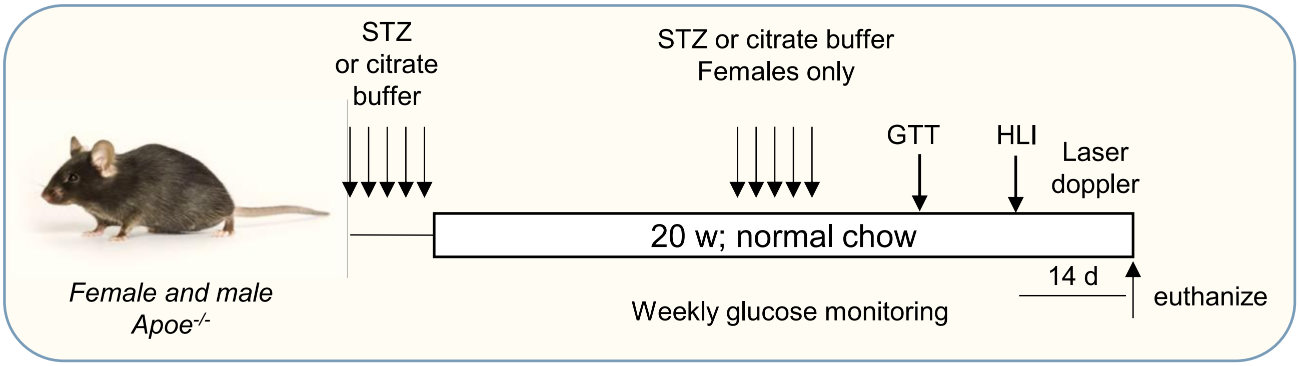


B.

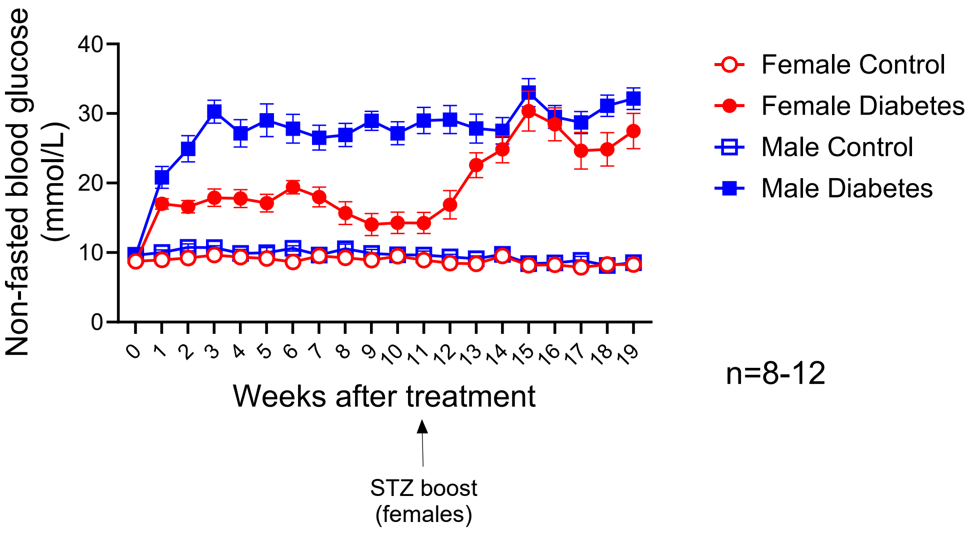


C.

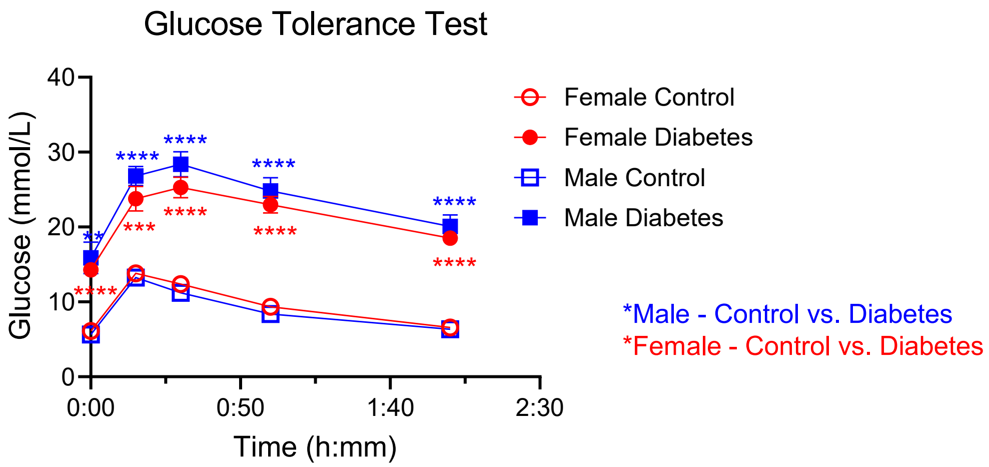

### Figure S2: Plaque size is unchanged with sex

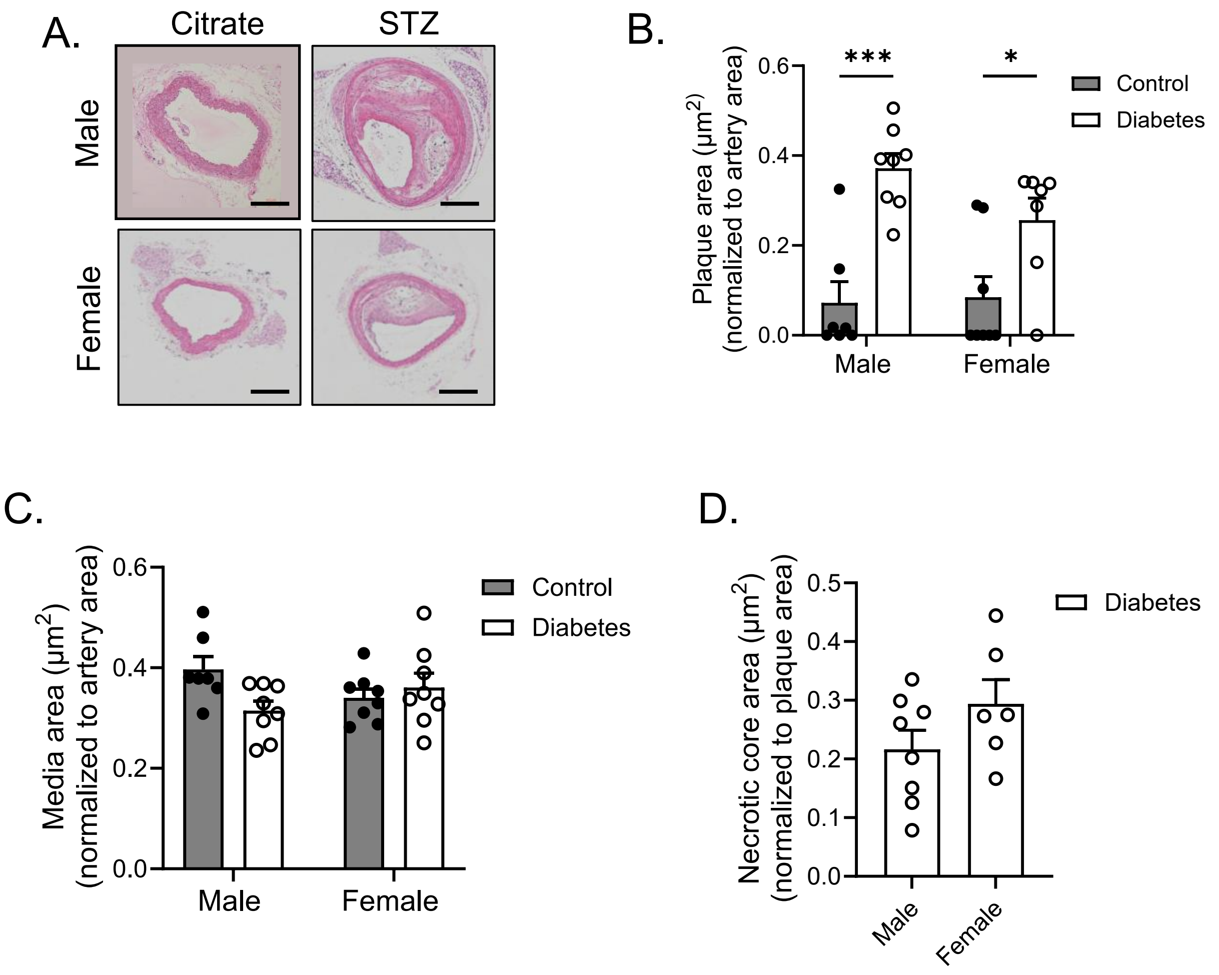

#### Figure S3: Female mice exhibit increased aortic oxidative stress and inflammation

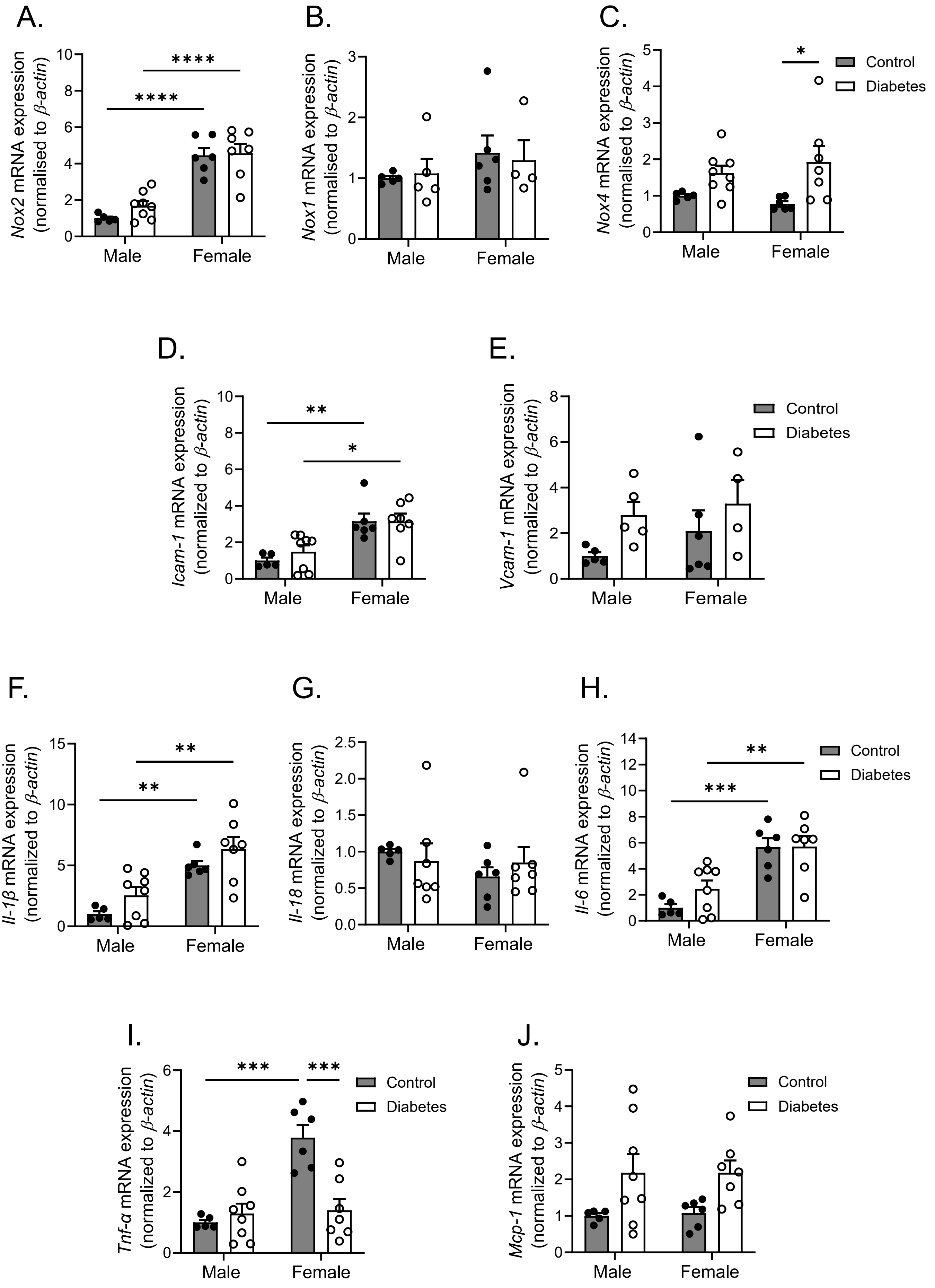

### Figure S4: Female ischemic limbs have increased oxidative stress and inflammatory gene changes

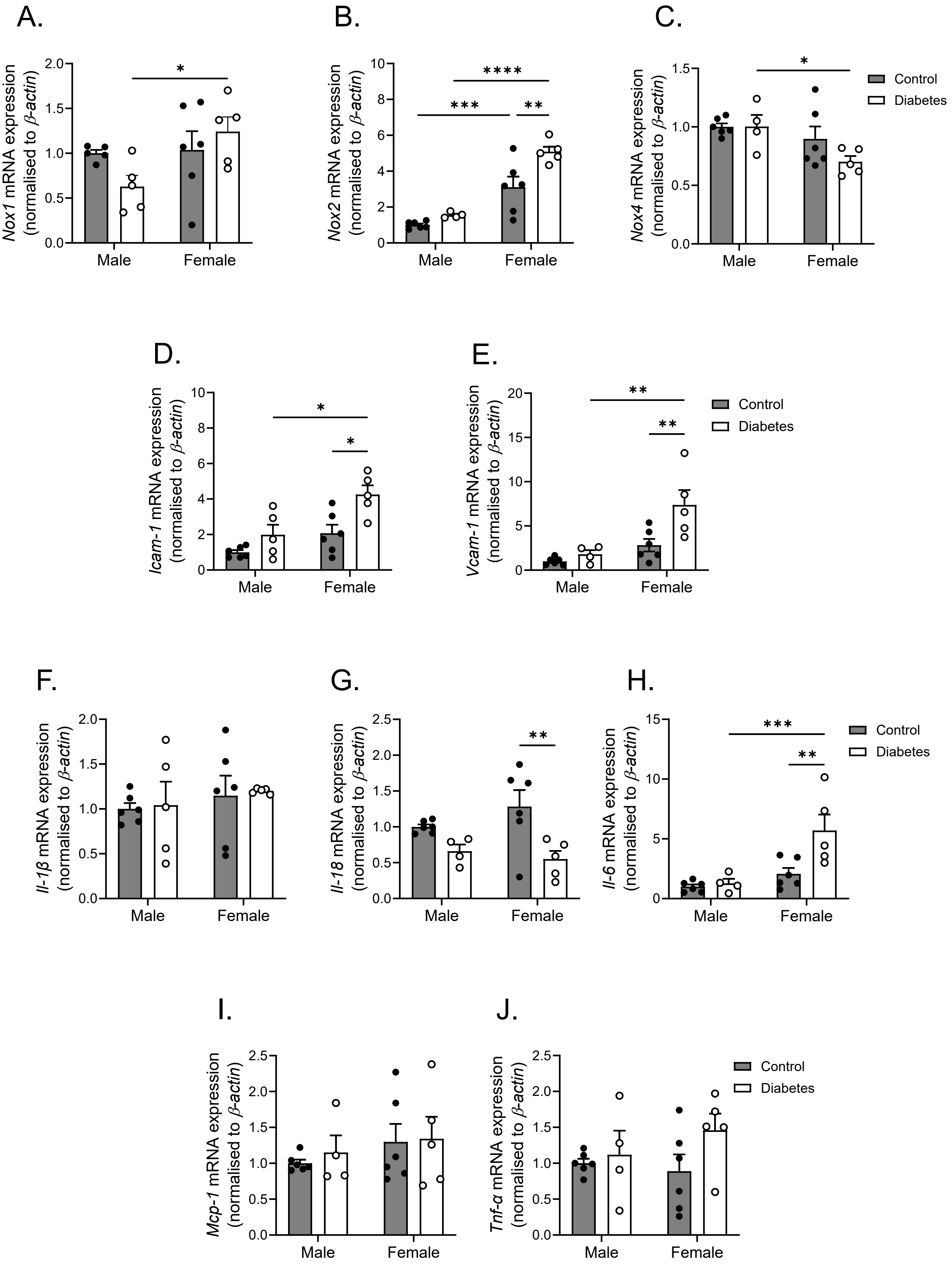

#### Figure S5: Angiogenesis and wound healing are reduced in female diabetic mice

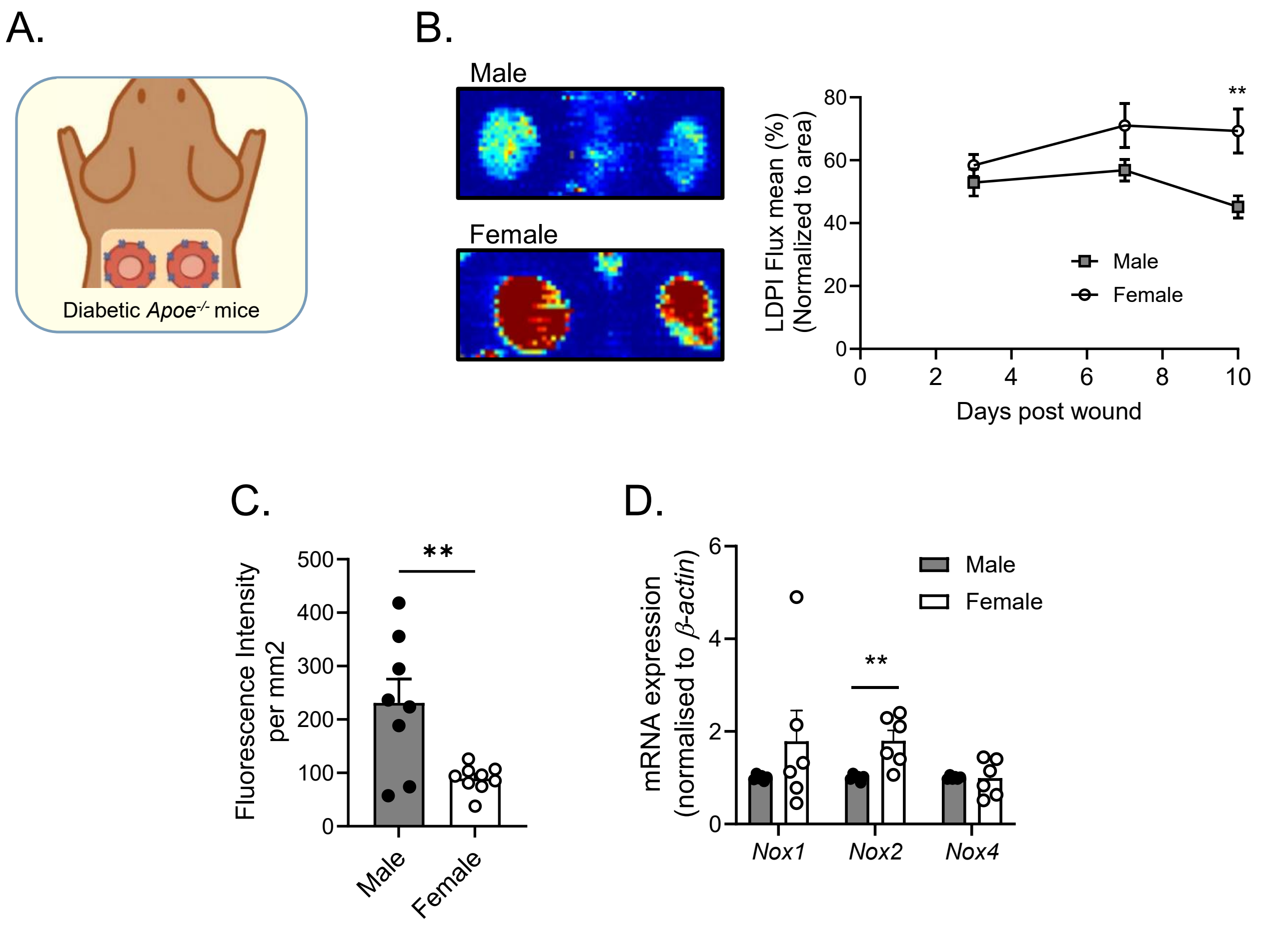
